# The ALS-associated co-chaperone DNAJC7 mediates neuroprotection against proteotoxic stress by modulating HSF1 activity

**DOI:** 10.1101/2024.12.01.626216

**Authors:** Andrew C. Fleming, Nalini R. Rao, Matthew Wright, Jeffrey N. Savas, Evangelos Kiskinis

## Abstract

The degeneration of neurons in patients with amyotrophic lateral sclerosis (ALS) is commonly associated with accumulation of misfolded, insoluble proteins. Heat shock proteins (HSPs) are central regulators of protein homeostasis as they fold newly synthesized proteins and refold damaged proteins. Heterozygous loss-of-function mutations in the *DNAJC7* gene that encodes an HSP co-chaperone were recently identified as a cause for rare forms of ALS, yet the mechanisms underlying pathogenesis remain unclear. Using mass spectrometry, we found that the DNAJC7 interactome in human motor neurons (MNs) is enriched for RNA binding proteins (RBPs) and stress response chaperones. MNs generated from iPSCs with the ALS-associated mutation R156X in *DNAJC7* exhibit increased insolubility of its client RBP HNRNPU and associated RNA metabolism alterations. Additionally, DNAJC7 haploinsufficiency renders MNs increasingly susceptible to proteotoxic stress and cell death as a result of an ablated HSF1 stress response pathway. Critically, expression of HSF1 in mutant DNAJC7 MNs is sufficient to rescue their sensitivity to proteotoxic stress, while postmortem ALS patient cortical neurons exhibit a reduction in the expression of HSF1 pathway genes. Taken together, our work identifies DNAJC7 as a crucial mediator of HNRNPU function and stress response pathways in human MNs and highlights HSF1 as a therapeutic target in ALS.

## INTRODUCTION

Amyotrophic Lateral Sclerosis (ALS) is a devasting neurodegenerative disease which is driven by the dysfunction and degeneration of upper and lower motor neurons (MNs)^1,2^. The progressive loss of MNs leads to muscle atrophy and diminished ability to stimulate and control movement^1,2^. ALS cases are predominantly sporadic in nature, but a relatively small proportion of patients (<12%) suffers from familial forms of the disease^3^. The genetic etiology underlying these familial cases is complex, comprised of over 30 disease-causing genes encoding functionally diverse proteins involved in cellular functions ranging from RNA processing, cytoskeletal homeostasis, and axonal transport to protein quality control pathways^4–6^.

*DNAJC7* is a recently identified ALS-associated gene^7^ that is of particular interest because it encodes a co-chaperone protein with an ostensible but relatively unexplored role in protein homeostasis^8^, a process known to be perturbed in ALS and other neurodegenerative diseases^9^. It was first identified as an ALS-associated gene in a whole-exome sequencing (WES) study that highlighted a significant enrichment for protein truncating variants (PTVs) in *DNAJC7*, including nonsense and rare missense variants predicted to be pathogenic^7^. Importantly, additional disease-causative variants in *DNAJC7* have been identified in a series of subsequent independent studies on small cohorts of ALS patients^10–15^. Most ALS enriched variants in *DNAJC7* are nonsense, while many missense variants are located within functional protein domains, strongly suggesting disease mechanisms driven by DNAJC7 haploinsufficiency^16^. However, the mechanisms by which mutant DNAJC7 causes MN dysfunction and eventual degeneration remain unknown.

The DNAJC7 protein is highly expressed within the central nervous system (CNS) and belongs to a superfamily of co-chaperone proteins known as HSP40s that are responsible for maintenance of protein homeostasis processes including *de novo* protein folding and cellular stress response pathways^8,16,17^. The specific role of the HSP40s co-chaperones such as DNAJC7 is to facilitate the unambiguous selection of substrates for HSP70/HSP90 chaperones amongst the promiscuous pool of available client proteins^8,18^. As such, the known interacting partners of DNAJC7 include several protein components of the HSP family including HSP70s (e.g., HSPA4, HSPA8) and HSP90s, where it has been shown to “bridge” clients between HSP70 and 90^8,16–20^. While very little is known about the specific role or binding partners of DNAJC7 in neurons, the HSP70/90 complexes play critical roles in cellular proteostasis pathways. One such pathway is the heat shock response (HSR) pathway, which is governed by the highly conserved master transcription factor heat shock factor 1 (HSF1), and is broadly utilized by cells to cope with proteotoxic stress (e.g., stress caused by misfolded proteins) by upregulating the expression of heat shock proteins (HSPs) and chaperones^21–24^.

Here, we sought to determine how mutant DNAJC7 causes MN dysfunction. We discovered novel DNAJC7 interactors in human MNs, which were enriched for proteins participating in RNA metabolism and cellular responses to stress. Using CRISPR-Cas9 edited iPSC-derived MN models, we found that DNAJC7 haploinsufficiency disrupts mRNA metabolism through increased insolubility of HNRNPU. Most notably we found that mutant DNAJC7 MNs are sensitized to proteotoxic stress and exhibit degeneration because of a disruption in the timely activation of HSF1. Overexpression of HSF1 in mutant DNAJC7-MNs rescued their premature degeneration upon prolonged exposure to proteotoxic insult. Our findings elucidate the mechanisms by which DNAJC7 regulates MN proteostasis and highlight the potential of HSF1 stimulation as a therapeutic target in ALS to counter MN vulnerability.

## RESULTS

### DNAJC7 interacts with RNA binding proteins and regulators of stress response pathways in human MNs

In order to investigate the functional role of DNAJC7 in the context of ALS we sought to identify its binding partners in human MNs, which represent the most vulnerable cell type in the disease. We used a healthy control iPSC line (line CS0002, see Methods) to generate lower MNs^25^, and performed immunoprecipitation (IP) of endogenous DNAJC7, followed by liquid chromatography tandem mass spectrometry (LC-MS/MS)-based proteomic analysis (Figure 1A-B and S1A). We conducted the IP-MS experiment using MN cultures from 3 independent differentiations and identified 88 proteins that were consistently co-purified within the DNAJC7 IP (Figure 1C). Of these, 48 proteins were found to co-purify exclusively with DNAJC7, while 13 were significantly enriched within the DNAJC7 IPs, relative to the IgG control IPs (p<0.05) (Figure 1C, Figure S1B). To add further context to the DNAJC7 interactome we performed STRING and gene ontology (GO) bioinformatic analysis, which collectively highlighted “RNA binding”, “ATP binding”, “cellular response to stress” and “cytoskeleton” as significantly enriched categories (Figure 1D-E and S1C). Importantly, these terms have been previously implicated in ALS pathogenesis in both genetic analysis and observations from disease models and post-mortem patient tissue^26–29^. We specifically identified several RNA-binding proteins (RBPs) including MATR3 and members of the family of heterogeneous ribonucleoprotein particle (HNRNP) proteins such as HNRNPK, HNRNPU, HNRNPD, HNRNPC and HNRNPA1 (Figure 1C and Table S1). Notably, genetic mutations in *MATR3* and *HNRNPA1* themselves are responsible for rare forms of ALS^30^, suggesting a converging functional interaction between independent causal genes. DNAJC7 has been previously shown to interact with another ALS-causal RBP in FUS^31^, although we did not confirm this in our unbiased proteomics analysis. The “cellular response to stress” enrichment was driven by several HSPs with chaperone activity including HSPA1A, HSPA8, and HSP90 (Figure 1C and Table S1). In fact, one of the main DNAJC7 interactors was HSPA1A, an HSP70 that is tightly linked to stress response pathways and has previously been shown to interact with DNAJC7 in multiple non neuronal cell models^8,18,19,32^. To independently validate these findings, we performed IP-Western blot (WB) analysis and probed for representative proteins from the “stress response” and “RNA binding” pathways in HSPA1A, HSP90AB1 and MATR3 respectively (Figure 1F-H). Collectively, analysis of the DNAJC7 interactome in MNs showed that it physically interacts, potentially through HSP70/90 complexes, with a small and specific set of proteins enriched for RBPs and HSPs.

**Figure 1.**
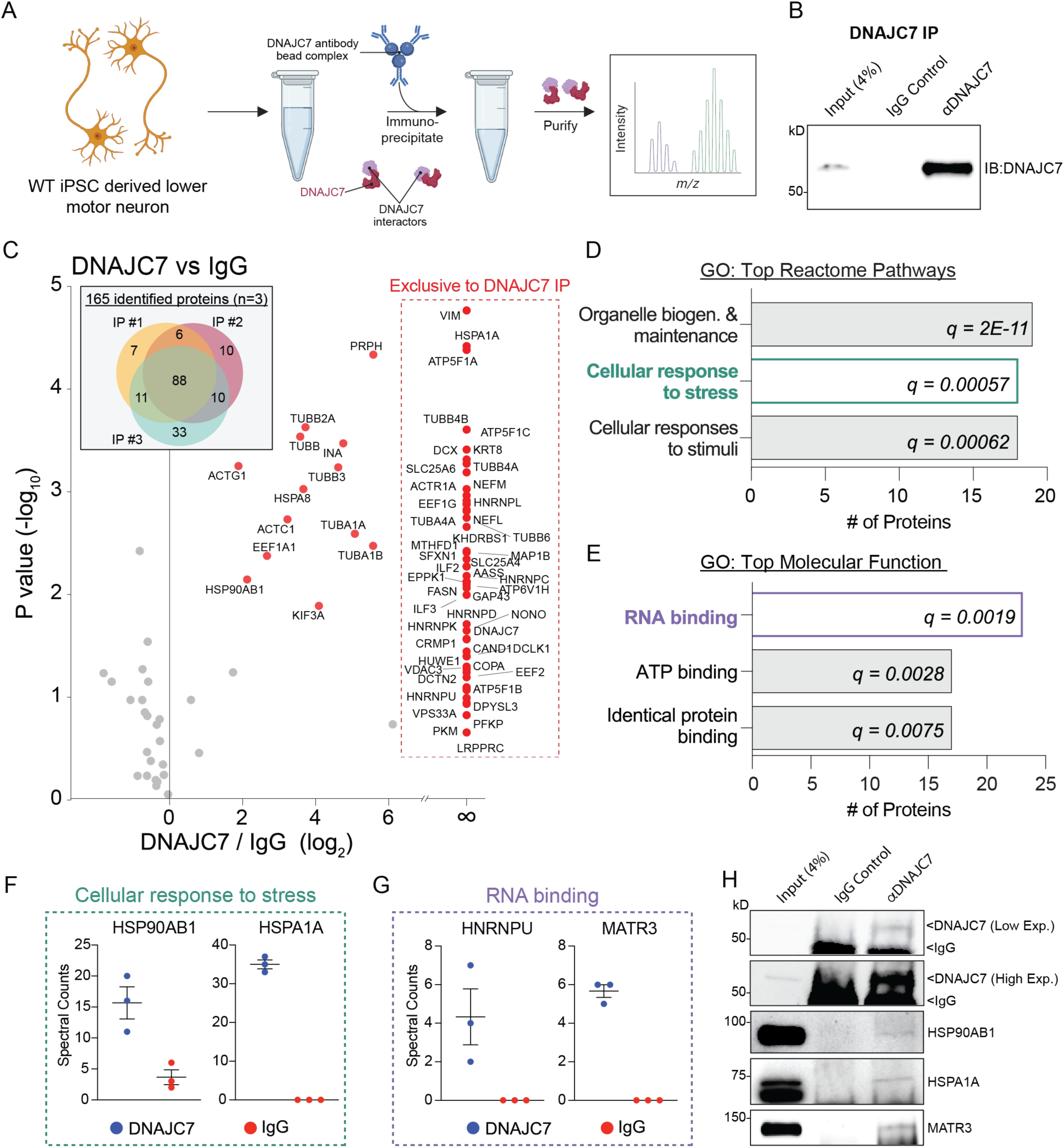
DNAJC7 interacts with RNA binding proteins and regulators of stress response pathways in human MNs. (A) Schematic of co immunoprecipitation of DNAJC7 from WT MNs. (B) WB of DNAJC7 from co-IP. Input = 4% of total protein used in co immunoprecipitation. (C) Venn diagram: all identified proteins from each DNAJC7 IP (n=3), 88 of which persistently identified proteins from each biological replicate. Scatter plot: Gray dots represent proteins not significantly enriched in DNAJC7 IP, red dots represent proteins significantly enriched in DNAJC7 IP fraction (p<0.05) or found exclusive in DNAJC7 IP (61 proteins). (D and E) Gene ontology of pathways overrepresented within DNAJC7 interactome, filtered by fold enrichment and ranked by number of proteins. FDR = q value. (F and G) Plot of spectral counts of proteins found in DNAJC7 IP vs IgG IP. Values represent the mean ± standard error of the mean (SEM). Each dot represents a biological replicate (N = 3). (H) WB of proteins co immunoprecipitated with DNAJC7. Input = 4% of total protein used in co immunoprecipitation. DNAJC7 WB: top band = DNAJC7 protein, lower band = IgG heavy chain visualized at low and high exposure. Note that IgG heavy chain does not appear in panel B because the DNAJC7 antibody was covalently linked to immunoprecipitation beads.

### DNAJC7 haploinsufficiency disrupts HNRNPU solubility and target mRNA expression

We next sought to develop a cellular model to investigate the functional ramifications of ALS-associated DNAJC7 mutations (Figure 2A). We used CRISPR-Cas9 to knock-in a premature truncation point mutation (p466 C>T [R156X]) into the endogenous gene locus in a healthy control iPSC stem cell line (line CS0002, see Methods) (Figure 2B). We targeted this mutation out of more than 15 that have been reported so far (Figure 2A) because is the most common ALS-causing variant^7,16^, and it is predicted to be deleterious. The editing generated an isogenic pair of iPSC clones with two distinct genotypes: a heterozygous clone containing one copy of the mutation (DNAJC7^R156X/+^), and a targeted but unedited isogenic wild type (WT) control (Figure 2B). WB analysis showed that the DNAJC7 protein was reduced by ∼75% in heterozygous mutant iPSC-derived MNs (p<0.0001), consistent with previous work demonstrating a similar level of DNAJC7 reduction in ALS patients harboring 1 copy of R156X^7^ (Figure 2C). Importantly, DNAJC7 haploinsufficiency did not impede the efficiency of motor neurogenesis, as the mutant DNAJC7 and isogenic control iPSC lines exhibited equal efficiency of differentiation as measured by immunofluorescence (IF) for the MN markers ISL1/2 and MAP2 (Figure S2A-C).

**Figure 2.**
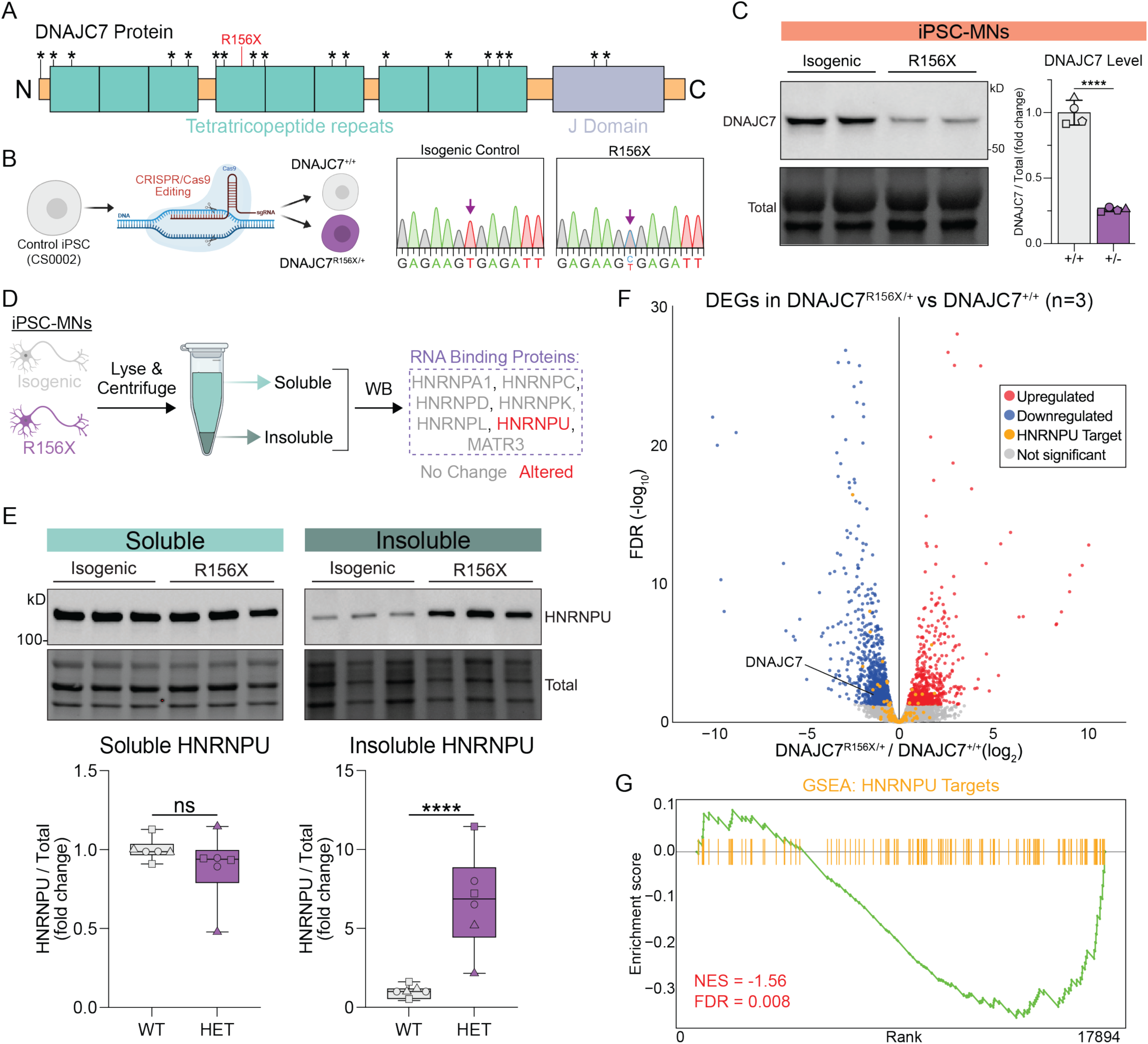
DNAJC7 haploinsufficiency disrupts HNRNPU solubility and target mRNA expression. (A) Schematic of known ALS causative mutations on DNAJC7 protein marked with * (B) Schematic of CRISPR-Cas9 knock-in of R156X mutation into healthy control iPSC line. (C) Left: WB images of DNAJC7 proteins levels from MN lysate derived from isogenic pairs (+/+ = isogenic control, +/- = R156x mutant). Bottom: Values represent the mean ± standard error of the mean (SEM). Four experiments are represented by distinct shaped symbols. N = 4, Unpaired t test (two-tailed): **** P<0.0001. (D) Schematic of biochemical fractionation of soluble/insoluble material from MN lysates followed by WB analysis. Red labels represent proteins whose solubility change in R156X mutant MNs. (E) Top: WB images of soluble and insoluble HNRNPU proteins levels from MN lysate derived from isogenic pairs. Bottom: Values represent the mean ± min/max. Experiments are represented by distinct shaped symbols. N = 6, Unpaired t test (two-tailed): **** P<0.0001. (F) Volcano plot of RNA Sequencing data from R156X vs control. Red or blue colored dots represent transcript significantly upregulated or downregulated, respectively (n = 3, FDR < 0.05). Yellow dots represent transcriptional targets of HNRNPU. (G) Gene set enrichment of HNRNPU eCLIP targets. Kolmogorov–Smirnov test: N = 3, NES = -1.56, FDR = 0.008.

Given the potential role of DNAJC7 in *de novo* protein folding, we next sought to utilize this isogenic iPSC platform to determine if DNAJC7 client RBPs have increased insolubility, indicating potential misfolding and subsequent dysfunction (Figure 2D). Intriguingly, we found that while most RBPs were not affected, the level of HNRNPU within the insoluble biochemical fraction, was significantly higher in mutant DNAJC7 MNs relative to isogenic controls (p<0.0001) (Figure 2E). Given HNRNPU’s relatively promiscuous role as an RNA binding protein, we next performed RNA-Sequencing (RNA Seq) in differentiated MNs to capture any potential HNRNPU- dependent changes in gene expression downstream of this altered solubility. We found ∼1600 differentially expressed genes in mutant DNAJC7 MNs (FDR < 0.05), and upon performing gene set enrichment analysis (GSEA) on known HNRNPU target mRNAs within the dataset, we found that they were significantly under- represented in DNAJC7^R156X/+^ cultures (normalized enrichment score [NES] = -1.56, FDR = 0.008) (Figure 2F- G, Figure S2D). As a negative control, we also performed GSEA on mRNAs associated with other RBPs (e.g., MATR3, HNRNPL) whose solubility was unaffected in mutant DNAJC7 MNs and found no significant enrichment (Figure S2E-G). Together, these data demonstrate that ALS-associated DNAJC7 haploinsufficiency impacts the solubility of HNRNPU and the expression level of some of its client mRNAs.

### DNAJC7 haploinsufficiency sensitizes MNs to proteotoxic stress

We next focused on dissecting the significance of DNAJC7 haploinsufficiency in cellular stress response pathways. Several chaperones including HSPA1A, HSPA8, and HSP90 that we found to bind DNAJC7 in MNs are involved in the handling of proteotoxic stress (Figure 3A). Thus, we hypothesized that loss of DNAJC7 might disrupt the ability of MNs to respond to proteotoxic stress and make them more sensitive to various relevant insults (Figure 3A). To test this hypothesis, we treated DNAJC7^R156X/+^ and isogenic control MNs with a series of small molecules that trigger endoplasmic reticulum (ER) stress (Brefeldin, 100nM), proteasomal stress (MG132, 5μM) and cytoplasmic stress (Ganetespib, 150nM) and monitored neuronal survival by live cell imaging (Figure 3B). To track the survival of individual cells we used a neuron-specific fluorescent reporter (*SYN1*::GFP) in combination with the cell death indicating dye propidium iodide (PI) (Figure 3B) and acquired single-cell resolution images of labeled MNs every 6 hours for 4 days after treatment (Figure 3C). We found that mutant DNAJC7 MNs degenerated significantly faster compared to their isogenic controls when exposed to MG132 (p<0.0001) and Ganetespib (p=0.0001), but not Brefeldin (p=0.8095), suggesting this increased sensitivity was relatively specific (Figure 3D-F). Importantly these effects were robust across n=3 independent differentiations (Figure S3A-C), while we did not observe any spontaneous degeneration after vehicle DMSO treatment in either genotype. Although the specific mechanism of action of MG132 and Ganetespib is different (MG132 blocks the proteosome and Ganetespib inhibits HSP90 activity), a notable converging consequence of both molecules is an accumulation of excess proteins^33–35^, suggesting that DNAJC7 haploinsufficiency renders MNs vulnerable to the toxic accumulation of proteins.

**Figure 3.**
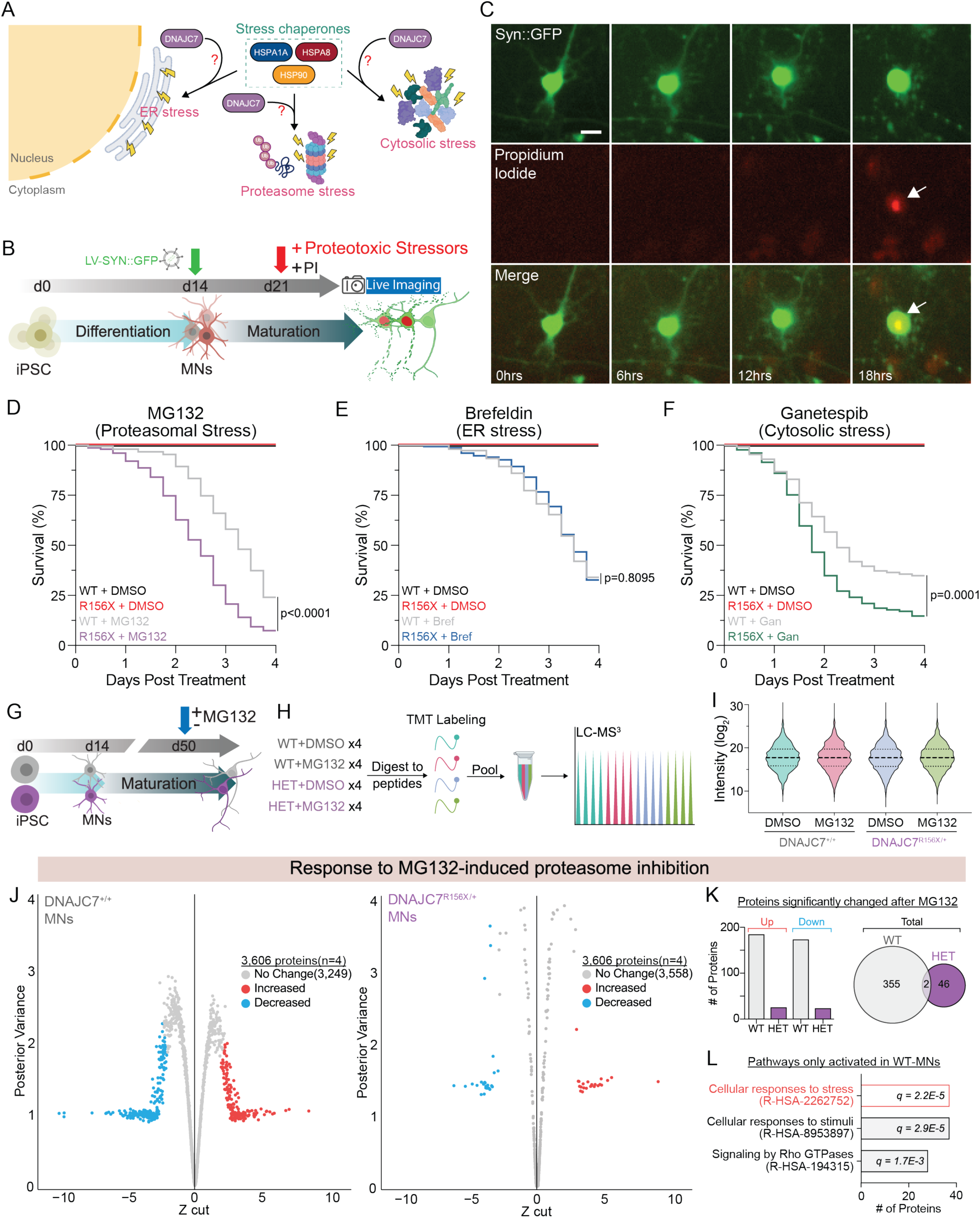
DNAJC7 haploinsufficiency sensitizes MNs to proteotoxic stress. (A) Schematic of stress pathways DNAJC7 may influence. (B) Schematic of live imaging experiments following proteotoxic stress induction. (C) Representative images of a dying MN co-labeled with SYN1-GFP (green) and propidium iodide (red) and merged. Scale bar, 20 μm. (D) Kaplan-Meier survival curve of MNs survival following MG132 or DMSO control. 150 cells tracked per condition, Mantel-Cox log-rank test: p<0.0001. (E) Kaplan-Meier survival curve of MNs survival following Brefeldin or DMSO control. 150 cells tracked per condition, Mantel-Cox log-rank test: p=0.8085. (F) Kaplan-Meier survival curve of MNs survival following Ganetespib or DMSO control. 129 cells tracked per condition, Mantel-Cox log-rank test: p=0.0001. (G and H) Schematic of MG132 treatment in MNs followed by tandem mass tag quantitative proteomics. (I) Violin plot of the average TMT intensities of all proteins for each condition. Two-way ANOVA: no significance (J) Shrinkage plots of protein level changes in MNs following MG132 treatment for R156X-MNs (right) and isogenic controls (left). Red and blue dots represent significantly increased or decreased proteins, respectively. N = 4 independent differentiations. (K) Left: bar graph displaying number of proteins changed in response to MG132 for each genotype. Right: Venn diagram showing lack of overlap of altered proteins. (L) GO enrichment of top reactome pathways upregulated in isogenic MNs following MG132 ranked based on # of genes in pathway. FDR = q value indicated in each bar.

We next used tandem mass tag mass spectrometry (TMT-MS) to compare the proteomic response of heterozygous DNAJC7^R156X/+^ and isogenic control MNs following 8 hours of MG132 treatment, as this stressor elicited a very robust survival phenotype (Figure 3G). TMT-MS employs multiplexed isobaric labeling that confers the advantage of enabling the pooling of all biological replicates, thereby eliminating batch variability and facilitating a highly quantitative method by which to directly compare the relative abundance of proteins from the same pool of identified peptides (Figure 3H). We quantified 3,606 proteins and critically found that the labeling efficiency was equal across all biological replicates and experimental conditions (Figure 3I). While most proteins were not alerted 8 hours after blocking the proteosome, 357 proteins, corresponding to approximately 9.9% of the quantified proteome, exhibited a significant change in their abundance in control MNs (Figure 3J-K and Figure S3D). In stark contrast, only 48 proteins or only 1.3% of the quantified proteome, exhibited measurable changes in mutant DNAJC7 MNs subjected to the same treatment (Figure 3J-K and Figure S3D). Notably, the proteins that become elevated in control MNs are enriched for pathways associated with “cellular responses to stress” suggesting a lack of such a canonical and potentially protective response in mutant MNs (Figure 3L). This robust lack of proteome remodeling in DNAJC7^R156X/+^ MNs is in accordance with their increased rates of degeneration in response to proteotoxic stress.

### DNAJC7 haploinsufficiency confers a basal reduction in HSF1 signaling in MNs

We next sought to better understand the molecular mechanisms driving the increased sensitivity of mutant DNAJC7 MNs to proteotoxic stress. We first carefully examined baseline gene and protein expression differences between DNAJC7^R156X/+^ and isogenic control MNs with a focus on stress-associated terms. We identified 398 significantly altered proteins between the two genotypes and observed that several of the proteins that were significantly downregulated in DNAJC7^R156X/+^ MNs represented known targets of the heat shock transcription factor 1 (HSF1) (Figure 4A). Accordingly, a targeted gene set enrichment analysis (GSEA) of 47 unique terms directly related to “stress”, revealed a highly significant downregulation of HSF1 target proteins (FDR<0.0001) (MSigDB#: M19734) (Figure 4B-C). Critically, this effect was highly specific, as no terms associated with other types of stress such as DNA damage response (MSigDB#: M13636), unfolded protein response (MSigDB#: M5922), cellular response to oxidative stress (MSigDB#: M45123), or cellular response to chemical stress (MSigDB#: M29264) were significantly changed in mutant DNAJC7 MNs (Figure S4A). Two of the most downregulated HSF1 target proteins in the MS dataset were CRYZ (p=0.0002) and HSPB1 (p<0.0001) (Figure 4A), and we validated the reduction in their levels by WB in independently differentiated MN samples from DNAJC7^R156X/+^ and isogenic control iPSC lines (Figure 4D). GSEA on RNA-Seq datasets revealed a moderate but significant downregulation in the expression of HSF1 target genes in mutant DNAJC7 MNs (FDR = 0.02) (Figure S4B), further validating the effects on this pathway at both the transcriptional and proteomic level.

**Figure 4.**
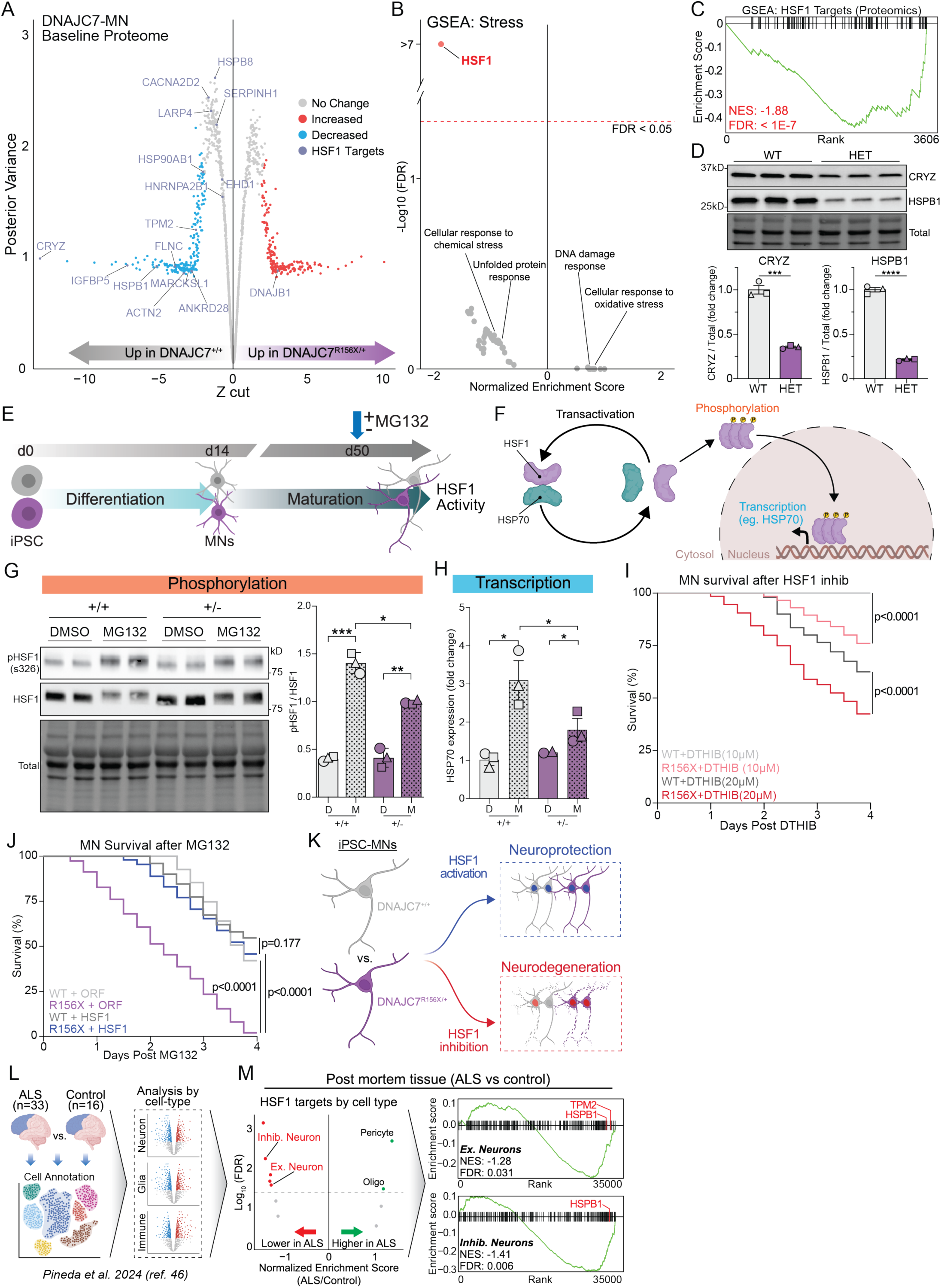
DNAJC7 regulates stress-induced activation of HSF1 and activation of HSF1 rescues the sensitivity of mutant DNAJC7-MN to stress. (A) Shrinkage plot of basal relative protein abundance between DNAJC7-MNs and isogenic controls. Purple dots represent HSF1 transcriptional targets. (B) Scatter plot of gene set enrichment analysis of “stress” terms. Colored or gray dots represent significantly (FDR < 0.05) or not significantly (FDR > 0.05) enriched terms, respectively. (C) GSEA plot of HSF1 targets within the basal proteomic dataset. Kolmogorov–Smirnov test: NES = -1.88, FDR < 1E-7. (D) Top: WB image of CRYZ and HSPB1 proteins levels from MN lysate derived from isogenic pairs at baseline. Bottom: Values represent the mean ± standard error of the mean (SEM). Experiments are represented by distinct shaped symbols. N = 3, Unpaired t test (two-tailed): CRYZ ***P=0.0002, HSPB1 ****P<0.0001. (E) Schematic of HSF1 activity assessment in MNs following MG132. (F) Schematic of HSF1 activation pathway. (G) Left: WB image of phospho-HSF1 (Ser326) and total HSF1 protein levels in MNs with and without MG132. Right: Values represent ratio of pHSF1/total HSF1 ± standard error of the mean (SEM). Experiments are represented by distinct shaped symbols. N = 3, Two-way ANOVA: *p<0.05, **p<0.01, ***p<0.001. (H) RT-qPCR quantification of HSP70 transcript levels, fold change relative to WT + DMSO control. Experiments are represented by distinct shaped symbols. N = 2-3, Two-way ANOVA: *p<0.05. (I) Kaplan-Meier survival curve of MNs survival following 10 μM or 20 μM DTHIB. 144 cells tracked per condition, Mantel-Cox log-rank test: WT vs R156X (10 μM) p<0.0001, WT vs R156X (20 μM) p<0.0001. (J) Kaplan-Meier survival curve of MNs survival following MG132 with ORF-LV or HSF1-LV. 150 cells tracked per condition, Mantel-Cox log-rank test: WT+ORF vs R156X+ORF p<0.0001, R156X+HSF1 vs R156X+ORF p<0.0001, R156X+HSF1 vs WT+HSF1 p=0.177. (K) Schematic of DNAJC7-MN increased stress resistance following HSF1 activation and selective degeneration after HSF1 inhibition. (L) Schematic of single cell RNA-Seq dataset from Pineda et al.,^46^ used for analysis. (M) Left: scatter plot of gene set enrichment analysis of HSF1 Targets (MSigDB#: M19734) by cell type. Each dot represents a different cell type. Colored or gray dots represent significant (FDR < 0.05) or not significant (FDR > 0.05) in ALS vs Control samples. Inhib = Inhibitory, Ex = Excitatory, Oligo = Oligodendrocyte. Right: GSEA plots of HSF1 Targets (MSigDB#: M19734) in ALS vs Control in excitatory neurons or inhibitory. Labeled ticks indicate specific HSF1 targets also significantly reduced in DNAJC7-MNs. Kolmogorov–Smirnov test: Ex. (NES = -1.28, FDR = 0.031), Inhib. (NES = -1.41, FDR = 0.006).

HSF1 is a highly conserved transcription factor that acts as a master regulator of cellular stress response, by activating a transcriptional cascade of chaperones, foldases and metabolic genes to counter proteotoxic stress^21–24^. Thus, a reduction in HSF1 signaling in heterozygous DNAJC7^R156X/+^ MNs may be central to their increased vulnerability to proteotoxicity. Notably, HSPA1A and HSPA8, two proteins which we found to interact with DNAJC7 in MNs, are known to play a direct role in regulating HSF1 activity^24,36,37^. The activation of this pathway has been previously associated with ALS pathophysiology, which is characterized by the accumulation of misfolded and aggregated proteins such as SOD1, FUS and TDP-43^38–43^. To add further context to the significance of HSF1 signaling for neuronal homeostasis we used publicly available single-cell RNA Seq datasets^44,45^ to examine the expression of HSF1 and its target mRNAs within distinct cell types in the human CNS (Figure S4C-D). We found that all neurons in the cortex and spinal cord including spinal MNs, expressed higher levels of HSF1 targets relative to glia, vascular and immune cells within the CNS, suggesting a potential reliance of this pathway for neuronal homeostasis.

### DNAJC7 haploinsufficiency delays stress-induced activation of HSF1

Given the converging evidence pointing to reduced baseline HSF1 signaling, we next interrogated the molecular activation of this pathway in mutant DNAJC7 and isogenic control MNs in response to stress (Figure 4E). Activation of the HSF1 pathway consists of several well-defined steps including a dynamic coupling/decoupling of HSF1 with HSP70, HSF1 hyperphosphorylation, nuclear translocation and ultimately transcriptional upregulation of stress-response target genes (Figure 4F). We first measured the level of phosphorylated HSF1 (Ser326), a marker of active HSF1, and found that while both sets of MNs showed increased phosphorylation after MG132 treatment, the ratio of pHSF1 over total HSF1 was significantly reduced in DNAJC7^R156X/+^ MNs (n=3; p=0.033) (Figure 4G). To directly measure stress-dependent HSF1 activity we used RT-qPCR to evaluate the transcriptional level of the canonical HSF1 target *HSP70* and found reduced upregulation in mutant DNAJC7 MNs (n=3; p=0.0107) (Figure 4H). These data suggest that DNAJC7 haploinsufficiency in MNs disrupts the timely activation of HSF1 upon induction of proteotoxic stress.

### Activation of HSF1 rescues the sensitivity of mutant DNAJC7-MN to stress

To directly determine the contribution of HSF1 signaling to the vulnerability of mutant DNAJC7 MNs we next evaluated the effect of inhibiting or activating this pathway on neuronal survival. We first treated mutant and wildtype MNs with the highly specific HSF1 inhibitor DTHIB, which acts by binding to the DNA-binding domain of HSF1 and performed longitudinal live cell imaging analysis as described before. At low doses of treatment (10μM), DNAJC7^R156X/+^ MNs demonstrated a ∼25% reduction in survival, whereas isogenic control MNs were virtually unaffected with only <1% reduction in survival (n=3; p<0.0001) (Figure 4I). Similarly, treatment with a higher dose of DTHIB (20μM) caused a substantially larger reduction in the survival of DNAJC7^R156X/+^ MNs (61%) relative to its effect in control MNs (40%) (n=3; p<0.0001) (Figure 4I), demonstrating a dose-dependent sensitivity of MNs to this molecule, which is dramatically heightened by DNAJC7 haploinsufficiency.

We next used a lentiviral vector to infect MG132-treated MNs with HSF1 (HSF1-LV) or an empty vector as a control (ORF-LV). We ensured that the overexpression level of HSF1 was equal across both genotypes (Figure S4F), and that the empty vector did not have any effect on neuronal survival (Figure S4E). As we observed previously (Figure 3D-F), mutant DNAJC7 MNs were substantially more vulnerable to proteasomal inhibition than controls (WT-ORF *vs.* R156X-ORF; n=3, p<0.0001) (Figure 4J). Remarkably an only 1.5-fold level of HSF1 overexpression selectively improved the survival of mutant DNAJC7 MNs by 40% (R156X-ORF *vs.* R156X-HSF1; n=3, p<0.0001) (Figure 4J). The effect of HSF1 was minor in isogenic control MNs, resulting in no quantifiable differences in survival between the two groups of HSF1-treated MN genotypes (WT-HSF1 *vs.* R156X-HSF1; n=3, p=0.177) (Figure 4J). Taken together, our results demonstrate that mutant DNAJC7-MNs *in vitro* are particularly sensitive to HSF1 modulation, with overexpression being neuroprotective and inhibition causing neurodegeneration (Figure 4K).

Lastly, given that our findings revealed a bidirectional sensitivity of mutant DNAJC7 MNs to HSF1 modulation, we investigated whether the HSF1 pathway is relevant in the context of sporadic ALS disease. We used a single-cell RNA Seq dataset^46^ derived from human post-mortem cortical tissue comparing ALS patients (n=33, sporadic and mutant C9orf72) to age-matched healthy controls (n=16) (Figure 4L). We employed a “pseudo-bulk” analysis pipeline to measure cell-type-specific effects in ALS patient and control samples and found a significant reduction in the expression of HSF1 target genes in ALS inhibitory neurons (FDR = 0.006) and excitatory neurons (FDR = 0.031) relative to controls (Figure 4M, left). Moreover, targeted GSEA showed a significant downregulation of HSF1 pathway genes in cortical ALS neurons including HSPB1 (Figure 4M, right), analogous to the transcriptomic (Figure S4B) and proteomic (Figure 4C) profiles of iPSC-derived DNAJC7-MNs. This analysis suggests that HSF1-associated dysfunction may be a broadly pervasive feature of ALS.

## DISCUSSION

HSPs play a crucial role in the proteostasis deficits that pervade ALS pathophysiology. However, how loss of DNAJC7, which represents the only genetically linked HSP to ALS, causes MN dysfunction had remained unknown. To address this gap in knowledge, we identified DNAJC7 interacting partners and developed human iPSC-based models to interrogate functional defects in DNAJC7-deficient spinal MNs. We found that DNAJC7 haploinsufficiency reduced the solubility of the RBP HNRNPU and caused alterations in associated target mRNAs. We also found that DNAJC7 haploinsufficiency reduced the basal activity of HSF1 and impeded the timely activation of the HSF1 pathway in response to proteotoxic stress. These defects, which caused substantial vulnerability of MNs to proteotoxic stress, could be rescued by mild HSF1 overexpression. Our findings establish an indispensable role for HSF1 signaling in MN homeostasis, demonstrate that DNAJC7-associated sensitivity to stress is driven by a disruption in HSF1 activation, and highlight the therapeutic potential of targeting HSF1 to combat MN degeneration.

Intriguingly, we found a strong enrichment of RBPs that associate with DNAJC7 in human MNs including two proteins that can cause rare forms of ALS when mutated^30^, MATR3 and HNRNPA1, suggesting a converging interaction. We focused on HNRNPU because we found clear evidence for increased insolubility for this protein in mutant DNAJC7 MNs, at least under basal conditions. HNRNPU is an RBP that targets multiple RNAs to regulate their splicing and expression and plays a key role in cortical development^47^, while mutations in *HNRNPU* have been associated with rare forms of pediatric neurodevelopmental epilepsy^48^. The role if any at all, of HNRNPU in ALS pathophysiology has not been extensively explored, although it has been shown to directly bind to TDP-43 in nucleus and modulate TDP-43-dependent splicing and neurotoxicity^49^. It has also been flagged as an RBP with strong relevance to ALS by AI-based analysis of gene expression datasets and published literature^50^. While we found a disruption in the HNRNPU target gene expression signature in DNAJC7^R156X/+^ MNs, our work did not explore the functional relevance of these alterations or the potential implications of this phenotype in the context of sporadic ALS disease.

Beyond RBPs the DNAJC7 interactome analysis we conducted in MNs identified several components of cytoskeletal homeostasis including the microtubule proteins TUBA1B and TUBA4A. Although we did not observe any overt defects in relevant pathways such as neuronal morphology in our *in vitro* models of DNAJC7, cytoskeletal defects have been directly implicated in other forms of genetic ALS as well as sporadic disease^29,51,52^. Interestingly, recent studies have highlighted a functional interaction between DNAJC7, and the microtubule associated protein tau that is encoded by *MAPT*^53,54^. DnaJC7 was shown to coprecipitate with soluble human tau from brain preparations of a tauopathy mouse model and displayed higher binding affinity to natively folded WT relative to mutant tau, suppressing tau aggregation^53^. In a separate study, DNAJC7 was purified with insoluble mutant tau in HEK-293 cells, and subsequent knockdown of DNAJC7 both decreased the clearance of aggregated tau and exacerbated the seeding of tau *in vitro*^54^. Interestingly, overexpression of WT DNAJC7 abrogated mutant tau seeding, while introduction of ALS-associated mutations on the J-Domain of *DNAJC7*, which mediates interaction with HSP70, nullified this rescue. Tau aggregates are seen in as much as 50% of frontotemporal dementia (FTD) patients^55^, a neurological disorder characterized by speech and executive dysfunction that shares genetic and pathological overlap with ALS, while 50% of ALS patients are also characterized by FTD symptoms. Although we did not specifically identify tau in our native human MN models, these studies^53,54^ flag another potential neurodegenerative disease-relevant function of DNAJC7. Additionally, it is currently unknown whether ALS patients with *DNAJC7* mutations exhibit any tau pathology and whether mutations in *DNAJC7* can cause FTD-TAU. The continuous characterization of DNAJC7 patients and expanding ALS/FTD genetic studies will provide important clarity.

Our findings on DNAJC7 are in strong accordance with multiple studies that have demonstrated the neuronal protection qualities of other DNAJ co-chaperones in ALS, FTD, and Parkinson’s disease models, highlighting the converging importance of this class of proteins in neurodegenerative disease through their critical role in neuronal proteostasis^53,56–60^. In accordance with the canonical role of DNAJ co-chaperones we found that DNAJC7 was associated with several other HSPs, including HSP70s and HSP90s, which are involved in the modulation of cellular stress responses^36,37,61–63^. Consequently, we found DNAJC7-deficient MNs were highly sensitive to proteotoxic stress induced by small molecule treatments including MG132, which causes a buildup of proteins by blocking the proteasome, and Ganetespib that leads to a similar toxic accumulation of proteins by decoupling HSP90 from its clients^34,35^. These molecules undoubtedly trigger several stress pathways, but we focused on HSF1 based on unbiased gene and protein expression data supporting a DNAJC7-associated reduction in the levels of HSF1 associated chaperones both at basal conditions, as well as after stress induction. The HSR is a highly conserved proteostasis response and forms a cyclical regulatory network with several HSPs including DNAJC7 and its interacting partners such as HSP70 and HSP90^22,62^. HSF1 is a master transcription factor that elicits the HSR cascade by binding to heat shock elements (HSEs) on the promoter regions of heat shock responsive genes and activates their expression^21–23^. HSP70 and HSP90 are two such HSR genes, and upon sufficient protein expression act to suppress the activity of HSF1, forming a closed loop of autoregulation^36,37,61–63^. We provide experimental evidence supporting a key regulatory role for DNAJC7 in the timely activation of HSF1, likely through its co-chaperone activity on HSP70 and HSP90.

The canonical transactivation of the HSF1 pathway follows a cyclical HSP70-titration model and is thought to be driven by the dynamic coupling/uncoupling of the inhibitory HSP70/HSF1 complex^24^. This on/off chaperone switch is finely tuned^37^, and any disruptions in its cycling render the acute HSF1 stress response largely inadequate^36^. We found that iPSC-MNs show high expression levels of established HSF1 target genes such as HSPB1 and CRYZ, which were downregulated in mutant DNAJC7 MNs. These findings were also supported by our analysis of gene expression in CNS tissues that showed healthy control cortical neurons and spinal MNs distinctly favoring high HSF1 signaling. These data suggest that the pathway might already be active in MNs. We have similarly previously shown that iPSC-MNs as well as human spinal cord tissue neurons exhibit high basal levels of ER-stress as measured by XBP1-splicing^64^. It may be that post mitotic cells such as MNs constitutively utilize these stress pathways in order to cope with accumulated proteostasis insults over their especially long lifespan^65,66^. In this context, a DNAJC7-dependent disruption in HSF1 signaling would sensitize MNs to age-associated stress. At the same time, mouse MNs have previously been shown to maintain a high threshold of induction of the HSF1-mediated stress response relative to other cell types including glial cells^67^, with the suggestion that this contributes to their vulnerability to stress signals such as insoluble proteins. How exactly DNAJC7 depletion disrupts HSF1 activation remain to be determined but given its strong interaction with HSP70 and HSP90 it likely disrupts the HSF1 transactivation complex.

Our work highlights the HSF1 pathway as a therapeutic target in ALS patients with DNAJC7 mutations, and perhaps beyond^68^, as RNA-Seq analysis revealed a reduction in the expression of HSF1 pathway genes in sporadic and C9orf72 patient neurons. There have been several notable associations between ALS pathophysiology and dysfunction in elements directly downstream of HSF1 activation in response to stress, including accumulation of misfolded and aggregated proteins such as mutant FUS, SOD1 and OPTN^38–41^, perturbed HSP-mediated autophagic clearance of TDP-43^42,43^, and reduced expression of HSP40 and HSP70 in sporadic ALS spinal cord tissue^26^. Notably, HSF1 has been shown to prolong the survival of a mutant SOD1- ALS mouse model^69^, while *Chen et al*., found evidence of a compromised HSF1 response pathway in transgenic TDP-43 mouse models and ALS patients, and showed that a constitutively active form of HSF1 reduced the level of insoluble TDP-43 in mammalian cells^26^. Similarly, studies have shown that both FUS and C9ORF72 mutations trigger the HSF1 stress response pathway as an early means of neuronal protection, further illustrating the pervasive relevance of the timely activation of HSF1 in ALS^70,71^. The therapeutic potential of HSF1 is further bolstered by the finding that Riluzole, an FDA-approved drug that has a moderate but reproducible effect on extending ALS patient survival^72^, is thought to at least in part, elicit its benefit by increasing latent HSF1 levels^73^. At the same time, a phase 3 clinical trial in ALS patients using Arimoclomol^74^, which is a related HSP70 inducer that had shown promise in preclinical mouse models, was terminated as it did not lead to measurable efficacy outcomes^75,76^. HSF1 stimulation may be a challenging target with a narrow therapeutic index as most available molecules, agonize the pathway by causing mild stress^68^. A better understanding of the mechanisms by which HSF1 stimulation is protective in human MNs might allow for the design of more specific and safer molecules. Lastly, our findings underscore the importance of conducting a comprehensive characterization of this pathway and its specific gene targets in human MNs, which likely require a high basal level of HSF1 signaling for their homeostasis.

### Limitations of the study

While iPSC-models offer some unique advantages related to the ability of study human neurons, they are also limited on account of the fact that they represent *in vitro* models, without the context of an intact nervous system of a model organism. Additionally, our work is focused exclusively on the impact of DNAJC7 loss-of-function effects on spinal-like MNs, which are the cells that are primarily affected in ALS disease pathogenesis, but any potential non-cell autonomous effects remain unexplored. Lastly, while we show that HSF1 overexpression substantially ameliorates proteotoxic neurodegeneration of mutant DNAJC7 MNs, we did not define the mechanistic effects of HSF1 activation. It remains to be determined whether other genes that act downstream of HSF1 such the Bcl2-associated athanogene 3 (BAG3)^77^, a pro-survival co-chaperone to HSP70, would act in a similar way to protect mutant DNAJC7 MNs.

## ACKNOWLEDGMENTS

We are grateful to the following funding sources: US National Institutes of Health (NIH), National Institute on Neurological Disorders and Stroke (NINDS) and National Institute on Aging (NIA) R01NS104219 (E.K), NIH/NINDS grant R21NS131713 (E.K and J.N.S), S10 OD032464-01A1 (J.N.S.), the Les Turner ALS

Foundation (E.K) and the New York Stem Cell Foundation (E.K), NIH grant F31NS132580 (A.C.F). We would like to thank Marc Mendillo for helpful discussions and critical advice on the project. E.K is a Les Turner ALS Center Investigator and a New York Stem Cell Foundation – Robertson Investigator.

## METHODS

### Cell culture conditions

Induced pluripotent stem cells (iPSCs) were grown on Matrigel (Fisher), maintained with mTeSR1 medium (Stemcell technologies), and passaged every 5 days using Accutase (Innovative Cell Technologies). All cultures were kept at 37C with 5% CO2 and were grown in the absence of antibiotics. Furthermore, all cell lines used were tested monthly to ensure the absence of mycoplasma throughout the duration of this study.

### iPSC Models

Differentiation of lower motor neurons (MNs) from iPSCs were performed as previously described^29,78–80^. Briefly, ∼70% confluent iPSCs were dissociated using Accutate to achieve single-cell suspension and replated with mTeSR1 medium supplemented with 10µM ROCK inhibitor (Y-27632, DNSK International) at a density of 1.2 million cells/well of a 6 well plate. The next day, the mTeSR1 was removed and replaced with N2B27 differentiation medium (base of 50% Neurobasal and 50% DMEM:F12, supplemented with nonessential amino acids, GlutaMAX, N2, and B27; Gibco), further supplemented with small molecules: 10 μM SB431542 (DNSK International), 100 nM LDN-193189 (Tocris), 1 μM retinoic acid (RA; Sigma), and 1 μM Smoothened Agonist (SAG; DNSK International). This medium was maintained and changed daily until day 6 in culture, and subsequently replaced with N2B27 differentiation medium supplemented with 1 μM RA, 1 μM SAG, 5 μM DAPT (Tocris), and 4 μM SU5402 (DNSK International) to generate postmitotic spinal MNs. These cultures were then fed on a daily basis until day 14 in culture, and next were dissociated using TrpLE Express (Gibco) supplemented with DNase I (Worthington). MNs were then plated directly on Matrigel (BD Biosciences) coated cell culture plates and growth in NBM (base of neurobasal medium supplemented with nonessential amino acids, GlutaMAX, N2, B27, ascorbic acid (0.2 μg/ml, Sigma), brain-derived neurotrophic factor (BDNF), ciliary neurotrophic factor (CNTF), and glial cell line-derived neurotrophic factor (GDNF) (10 ng/ml, R&D Systems). For experiments requiring imaging or survival analysis, MNs were plated initially onto pre-coated Matrigel surfaces and allowed to attach for 24 hours, and the following day primary mouse glia cells (harvested from P0 mixed male and female pups of the CD1 strain as described previously^80^) were plated on top of MNs.

### Coimmunoprecipitation followed by mass spectrometry or WB

MNs lysates were collected in IP buffer (10mM Hepes [pH 7.6], 100mM NaCl, 1mM dithiothreitol, 10% glycerol, 1% sodium deoxycholate, 0.1% SDS, 1% Triton X-100, 1x protease inhibitor cocktail, and 1x phosphatase inhibitor cocktail. Insoluble material from cell extracts was then cleared by centrifugation, and protein concentrations were determined with a BCA kit (Pierce). Endogenous DNAJC7 was immunoprecipitated from 1mg of protein with anti-DNAJC7 antibody (Abcam, ab179830). IP of the antigen was conducted using Dynabeads Protein A magnetic beads (Invitrogen) following the manufacturer’s protocol. *For preparation of mass spectrometry (MS)*: eluted IP’d material was purified for compatibility to MS detection by briefly being run through precast polyacrylamide gel (Bio-Rad) to embed the material, followed by cutting out entire gel stack and subject to further processing for MS-based proteomics. *For preparation of western bot*: eluted proteins were separated by SDS-PAGE and subsequent transfer to nitrocellulose membrane (Bio-Rad). Membranes were then blocked in tris-buffered saline (TBS) + 0.1% Tween 20 (Bio-Rad) + 5% nonfat dry milk (LabScientific) and subject to overnight incubation at 4°C primary antibodies: DNAJC7 (1:1000, Abcam), HSP90 (1:500, Santa Cruz), MATR3 (1:10,000, Abcam), HSPA1A (1:1000, Novus), HNRNPU (1:500, Santa Cruz), and HNRNPK (1:000, Cell Signaling). All primary antibodies were diluted in TBS + 0.1% Tween + 5% nonfat dry milk. After multiple washes with TBS + 0.1% Tween, membranes were incubated with their corresponding secondary anti-mouse and anti-rabbit HRP-conjugated antibodies (1:5000, LI-COR Biotechnology). Membranes were then exposed to SuperSignal Pico chemiluminescent (Thermo Fisher Scientific) and imaged by ChemiDoc XRS+ system (Bio-Rad).

### MN survival tracking

For MN survival experiments, lentiviral *SYN1-GFP* (PZ196, Addgene) was added to MNs in suspension prior to seeding onto ImageLock 96 well plates (Sartorius) precoated with Matrigel. MNs were then plated in NBM supplemented with neurotrophic growth factors BDNF, CNTF, GDNF, and AA with SYN1-GFP virus for 24 hours before undergoing a full medium change to remove virus. In the case of lentiviral rescue experiments, LV- ORF/HSF1/BAG3 were co-transduced with SYN1-GFP at this time. Following the 24-hour incubation period, viral medium are removed and replaced with fresh medium and primary mouse glial cells. MN medium is then replenished every 2 days. After 7 days (21 days in culture), MNs are then treated individually with one of several variety of pharmacological agents including DMSO control (Sigma), MG132 (Calbiochem), Ganetespib (Mendillo Lab via Selleckchem), Brefeldin (Mendillo Lab via InvivoGen), Direct Targeted HSF1 InhiBitor (DTHIB, Mendillo Lab via Medchem) supplemented with cell death indicator propidium iodide (1:5000, Sigma). MN survival was then tracked with live imaging using IncuCyte S3 system (Sartorius) for green (GFP) and red (PI) signal emitted from cells. Image analysis was performed using Fiji software (U. S. National Institutes of Health, Bethesda, Maryland, USA), where cells were marked as dead when PI displays focal accumulation in the nucleus.

### Lentivirus production

To produce viruses (SYN1-GFP, LV-HSF1 [VectorBuilder], LV-ORF [VectorBuilder], LV-BAG3 [VectorBuilder]), HEK-293 cells were co-transfected with individual target lentivirus plasmids combined with packaging plasmids pMD2.G and psPAX2 vectors with HilyMax (Dojino Molecular Technologies). Virus was then collected from the cell medium 72 hours after transfection, filtered through a 0.22 μm PVDF syringe filter, and concentrated by centrifugation at 25,000g for 2 hours at 4°C. Concentrated virus was then resuspended in neurobasal medium (Gibco), aliquoted in appropriate volumes, and stored at -80°C.

### Co-immunoprecipiation mass spectrometry

*Mass spectrometry sample preparation:* Proteins were submitted as gel bands. Bands were destained by successive washes in acetonitrile and 100 mM ammonium bicarbonate. Disulfide bonds were reduced by incubating in 20 mM dithiothreitol (30 minutes at room temperature). Resulting free thiols were capped by incubating in 50 mM iodoacetamide (30 minutes at room temperature protected from light). Bands were then washed in 100 mM ammonium bicarbonate prior to addition of 2 µg of trypsin enzyme (Promega) followed by overnight incubation at 37 °C. The following day, the digestion was halted by acidification with 20 % formic acid and the peptide solution was dried in a vacuum centrifuge. The samples were re-suspended in 30 µl of 0.1 % formic acid, sonicated for 5 minutes to fully dissolve, then benchtop centrifuged. The supernatant was transferred to a glass vial and placed in the cooled auto-sampler rack.

*LC-MS/MS analysis*: Peptides were analyzed by LC-MS/MS using a Dionex UltiMate 3000 Rapid Separation LC system coupled to a linear ion trap—Orbitrap hybrid Elite mass spectrometer (Thermo Fisher Scientific, San Jose, CA). Four-microliter peptide samples were loaded onto the trap column, which was 150 μm × 3 cm in-house packed with 3 μm ReproSil-Pur beads (New Objective, Woburn, MA). The analytical column was a 75 μm × 10.5 cm PicoChip column packed with 3-μm ReproSil-Pur beads. Solvent A was 0.1 % aqueous formic acid and solvent B was 0.1 % formic acid in acetonitrile. Peptides were separated on a 120-minute analytical gradient from 5 % to 40 % solvent B at a flow rate of 300 nl/min. The mass spectrometer settings included positive data-dependent acquisition (DDA) mode, a 2.40 kV source voltage, and 275 °C capillary temperature. MS1 scans were acquired from 400 to 2000 m/z at 60,000 resolving power and 1E6 automatic gain control in the orbitrap. The top fifteen most abundant precursor ions in each MS1 scan were selected for fragmentation with an isolation width of 1 Da. Collision-induced dissociation (CID) at 35 % normalized collision energy in the ion trap was applied for fragmentation. Previously selected ions were dynamically excluded from re-selection for 60 seconds. A value of 3E5 was set for the MS2 automatic gain control.

*Mass spectrometry data analysis:* Raw files were converted to mgf format and analyzed using the Mascot search engine (Matrix Science, London, UK. version 2.7). MS/MS spectra were searched against the SwissProt human database (2021 version). All searches included carbamidomethyl cysteine as a fixed modification and oxidized methionine, deamidated asparagine and glutamine, and acetylated N-terminal as variable modifications. Three missed tryptic cleavages were allowed. Mass tolerances of 10 ppm (MS1 precursor) and 0.6 Da (MS2) were applied. Results were imported into Scaffold 5 software (Proteome Software, Portland, USA) for visualization.

### Tandem-mass-tag (TMT)-mass spectrometry

*TMT- MS Sample Preparation:* TMT-MS sample preparation was performed as previously described^66,81^. Briefly, 200 µg of whole cell extracts were methanol-chloroform precipitated. Extracted protein was resuspended in 6M guanidine in 100 mM TEAB and further reduced of disulfide bonds with DTT, followed by alkylation of cysteine residues with IAA. Proteins were then digested overnight at 37°C with 3 μg Trypsin/LysC (Promega). The digest was then acidified with formic acid and desalted (C18 HyperSep columns). Peptides were resuspended in 100mM TEAB and100μg for used for each respective isobaric TMT tag. After a 75 min incubation at RT, the reaction was quenched with 5% (v/v) hydroxylamine to 0.3%. Isobarically labeled samples were then combined 1:1:1:1:1:1:1:1:1:1:1:1:1:1:1:1 and subsequently desalted. The sample was then fractionated using high pH reversed-Phase columns (Pierce) and dried before reconstituted in LC-MS Buffer A (5% acetonitrile, 0.125% formic acid) for LC-MS/MS analysis.

*TMT-MS Data Collection*: TMT-MS analysis was performed as previously described^66,81^. Briefly, samples were resuspended in 20 µl Buffer A (5% acetonitrile, 0.125% formic acid) and 3µg of each fraction was loaded for LC-MS analysis. Orbitrap Fusion was used to generate MS data. The chromatographic run was performed with a 4h gradient as previously described^66,81^. In MS3, the top ten precursor peptides were selected for analysis were then fragmented using 65% HCD before orbitrap detection. A precursor selection range of 400–1200 m/z was chosen with mass range tolerance. An exclusion mass width was set to 18 ppm on the low and 5 ppm on the high. Isobaric tag loss exclusion was set to TMT reagent. Additional MS3 settings include an isolation window = 2, orbitrap resolution = 60 K, scan range = 120 – 500 m/z, AGC target = 6*105, max injection time = 120 ms, microscans = 1, and datatype = profile.

*TMT-MS Data Analysis and Quantification*: TMT-MS data analysis was performed as previously described ^66,81^. In brief, protein identification, TMT quantification, and analysis were performed with The Integrated Proteomics Pipeline-IP2 (Integrated Proteomics Applications, Inc., http://www.integratedproteomics.com/). Proteomic results were analyzed with ProLuCID, DTASelect2, Census, and QuantCompare. MS1, MS2, and MS3 spectrum raw files were extracted using RawExtract 1.9.9 software (http://fields.scripps.edu/downloads.php). Fully and half-tryptic peptide candidates were included in search space, all that fell within the mass tolerance window with no miscleavage constraint, assembled and filtered with DTASelect2 (ver. 2.1.3). Static modifications at 57.02146 C and 304.2071 K at N-term were included. Minimum peptide number was 2. The target-decoy strategy was used to verify peptide probabilities and false discovery ratios ^82^. Minimum peptide length of six was set for the process of each protein identification and each dataset included a 1% FDR rate at the protein level based on the target-decoy strategy and Isobaric labeling analysis was established with Census 2 with no intensity threshold applied.

### CRISPR/Cas9 editing

Healthy control iPSC line CS0002 was purchased from Cedar Sinai. iPSCs were edited by Applied StemCell Inc. (Milpitas, CA) and described in *Simkin et al*^83^. Briefly, one million iPSCs were electroporated with a mixture of sgRNA and Cas9 in a ribonucleoprotein format and ssODN. A small, presumably mixed population was then subjected to PCR and Sanger sequencing analysis. Once the heterogenous culture displayed sufficient repair with qualified HDR, the cells were then single-cell cloned. Individual colonies were picked after 2 weeks in culture and expanded. Positive clones were then further expanded and resequenced to confirm correct genotype prior to being cryopreserved and shipped. Furthermore, in previous work, *Simkin et al* performed extensive quality control on all iPSC clones utilized in this study including genomic DNA PCRs and Sanger sequencing, genomic integrity and pluripotency assays, analysis of off-target Cas9 sites, and quantitative genotyping PCR-based copy number assays^84^.

### Soluble/Insoluble fractionation followed by WB

Protocol is adapted and modified from *Tsioras et al*^39^. Briefly, MNs lysates were collected in RIPA buffer (50mM Tris (pH 7.4), 150mM NaCl, 0.5% sodium deoxycholate, 0.2% SDS, 1% Triton X-100, 1x protease inhibitor cocktail, and 1x phosphatase inhibitor cocktail), sonicated 3 x 3s, 40V output (QSonica, LLC), and centrifuged at 20,000g for 20 minutes. The supernatant, which is the RIPA-soluble fraction, was removed, while the RIPA-insoluble pellet remaining was further washed twice in RIPA lysis buffer with centrifugations of 20,000g for 30 minutes were performed between washes. Following the final wash, the insoluble pellet was resuspended in 2x Laemmli Sample Buffer (Bio-Rad) and sonicated 3 x 5s at 70V output. Both soluble and insoluble fractions were then boiled at 95 °C prior to loading for SDS-PAGE and WB analysis and probed for anti-HNRNPU (1:500, Santa Cruz), anti-HNRNPL (1:5000, Novus), and anti-MATR3 (1:10000, Abcam).

### RNA Extraction

Cells were harvested in 1mL TRIzol Reagent (Thermo Fisher Scientific) per 1 million cells. 0.2mL of chloroform was mixed with the samples for 3 minutes prior to being centrifuged at 12,000g at 4°C for 15 minutes. The RNA containing aqueous phase was then transferred to a new tube and 5 μg of Glycogen carrier (Thermo Fisher Scientific) was added to facilitate complete precipitation. Then, 0.5mL of isopropanol was added to the mixtures and left to incubate at room temperature for 25 minutes prior to being centrifuged for 10 minutes at 15,000g at 4°C.

### RNA Seq data processing

RNA libraries were sent to Novogene for QC, and library preparation (250∼300 bp insert strand nonspecific library with polyA enrichment). Libraries were sequenced using the Illumna NovoSeq platform, targeting at least 80M 150bp paired-end reads per sample Raw reads were trimmed using cutadapt (v3.4) to remove Illumina universal adapter sequences ( -aAGATCGGAAGAGCACACGTCTGAACTCCAGTCA,- AAGATCGGAAGAGCGTCGTGTAGGGAAAGAGTGT), trailing N bases (--trim-n), and bases with Phred score < 10 (--nextseq-trim=10), as well as any reads that were too short after trimming (-m 25). Trimmed reads were aligned to Gencode V38 (GRCh38.p13) with STAR (v.2.7.5a), using the flag –twopassMode Basic and default settings. Read counts were quantified at the gene-level using featureCounts (v2.0.1), specifying -s 0 for unstranded libraries.

### Differential Expression

Gene-level differential expression analysis was performed in R with DESeq2 (v1.38.3). Lowly expressed genes were filtered out before analysis – a gene must have had at least 0.5 counts per million reads in at least 3 samples in order to be retained. Significant genes were defined as having FDR<0.05 and log2(FoldChange)>=1.

### Gene ontology, GSEA, and STRING analysis

Gene ontology (GO) analysis was performed using The Database for Annotation, Visualization and Integrated Discovery (DAVID) ^85,86^. Top terms were filtered by a statistical and fold enrichment cutoff and GraphPad Prism 10 was used to visualize the enriched GO enrichment. Gene set enrichment analysis (GSEA) was performed using GSEA-MSigDB software (v4.3.3)^87,88^. Standard parameters were used in the query but briefly are as follows: stress terms gene set database (49 queries), 10,000 phenotype permutations, platform: MSigDB.v2024.1.Hs.chip, weighted Signal2Noise ranking, meandiv normalization mode, and Wald statistic values (stats) from DESeq2 values from differential expression were used as ranking values for the genes. Kolmogorov–Smirnov test used to compute normalized enrichment score and false discovery rate correction. For STRING analysis, physical subnetwork analysis, where thickness of line represents confidence (scaled 0 to 1) with manual color annotation overlaid to corresponding relevant GO terms.

### HSF1 target enrichment in single cell data sets

Processed single cell data sets from cortical and spinal tissues were downloaded from Synapse (syn45351388) and GEO (GSE190442), respectively. Author-provided counts and cell type annotations were used to create individual Seurat objects for cortex and spine data sets, and counts data were normalized using the *NormalizeData()* function. Following normalization, expression of HSF1 targets (from the HSF1_01 MSigDb gene set) at a per-cell were determined using the *AddModuleScore()* function. Module scores of HSF1 targets were then visualized for each cell type. For pseudobulk analysis in each cell type, counts were aggregated for individual patients using the Seurat function AggregateExpression. Aggregated counts for each cell type were provided to DESeq2 for differential expression analysis between ALS patients and healthy controls. For GSEA analysis, genes were ranked by their DESeq2 test statistic and analyzed for HSF1 target enrichment as described previously.

### Immunocytochemistry

Cells were fixed with 4% paraformaldehyde (PFA) and blocked for 1 hour in phosphate-buffered saline (PBS) supplemented with 10% normal donkey serum (Jackson ImmunoResearch) and 0.1% Triton X-100. Cells were then incubated overnight at 4°C with primary antibodies: ISL1/2(1:100, DSHB), MAP2 (Abcam, 1:5000), and HSF1 (ProteinTech, 1:200). Primary antibodies were removed and washed several times with PBS + 0.1% Triton before being incubated with Hoeschst 33342 (Invitrogen) and the appropriate secondary antibodies conjugated to Alexa Fluor 488, Alexa Fluor 555, or Alexa Fluor 647 fluorophores (1:500, Thermo Fisher Scientific) for 2 hours at room temperature. After several more washes in PBS + 0.1% Triton, cells were imaged directly in 24- well #1.5 glass bottom coated tissue culture plates (Cellvis) or coverslips were mounted with Fluoromount-G (Thermo Fisher Scientific).

### RT-qPCR

Following RNA extraction, first-strand cDNA was synthesized from 1-2 μg of DNase I (Invitrogen) treated RNA using SuperScript IV reverse transcriptase (Thermo Fisher Scientific) and oligo dT primers following manufacturer’s instructions. Synthesized cDNA was first diluted 1:10-1:20 and 2uL of diluted cDNA was used in each RT-PCR reaction performed with SYBR green (Thermo Fisher Scientific) on CFX system (Bio-Rad). PCR was performed under the following conditions: 95°C for 3 min, 40 cycles at 95°C for 10 sec and 60°C for 30 sec, and final step from 65°C to 95°C in increments of 0.5°C every 5 sec. All reactions were performed in duplicate. Average cycle of threshold (Ct) value of housekeeping gene GAPDH was subtracted from the Ct value of the gene of interest to obtain the ΔCt. Relative gene expression was then defined as the 2^−ΔCt^ (ΔΔCt) and normalized to each indicated sample control indicated in each experiment. Primer sequences used in this study: HSP70_F, ACCTTCGACGTGTCCATCCTGA; HSP70_R, TCCTCCACGAAGTGGTTCACCA; GAPDH_F, ACAACTTTGGTATCGTGGAAGG; GAPDH_R, GCCATCACGCCACAGTTTC.

## Statistical Analysis

We performed all statistical analysis using Prism 9 software (GraphPad) and Fiji ((https://imagej.nih.gov/ij/). All values in figures with error bars are presented as mean ± standard error of the mean (SEM). We classified an independent biological replicate as an independent iPSC differentiation performed on a different day. All numbers (*n*), significance values (*P* or *Q* value), and statistical test performed are specified either in the Results section or in each specific corresponding figure legend. To ensure that pooling of data in each experiment was appropriate, we first tested whether datasets variances were significantly different using Brown-Forsythe analysis of variance (ANOVA) test. Following this, we next used the D’Agostino-Pearson test to test whether sample data fit a Gaussian distribution. For experiments that included at least n = 3, we performed one-way ANOVA followed by Tukey’s post hoc for parametric test, and Kruskal-Wallis rank test with Dunn’s correction for multiple comparisons for nonparametric test). All survival experiments were performed at least three times, with equal numbers of neurons from each replicate being used for quantification shown. Statistical analysis was then performed using a two-sided Mantel-Cox log-rank test from 40-50 neurons per replicate per experiment. All TMT-based proteomics experiments were analyzed with Bayesian analysis of variance using BAMarray 3.0, a Java software package that implements the Bayesian ANOVA for microarray (BAM) algorithm. The BAM approach uses a special type of inferential regularization known as spike-and-slab shrinkage, which provides an optimal balance between total false detections (the total number of genes falsely identified as being differentially expressed) and total false nondetections (the total number of genes falsely identified as being nondifferentially expressed)^89^. Details statistics for each Figure described below:

- Figure 1: (C) n = 3 experiments were from 3 independent differentiations of either DNAJC7 IP or negative control IgG IP. (D) individual q values from all enriched terms are as follows: Reactome, from top to bottom: 2E-11, 0.00057, 0.00062; Molecular Function, from top to bottom: 0.0019, 0.0028, 0.0075.
- Figure 2: (C) n = 4 independent differentiations; values represent the mean ± standard error of the mean (SEM), unpaired t test (two-tailed): **** P<0.0001. (E) n = 6 differentiations; values represent the mean ± min/max. unpaired t test (two-tailed): ns = 0.276, **** P<0.0001.
- Figure 3: (D) 150 cells tracked per condition, Mantel-Cox log-rank test: p<0.0001. (E) 129 cells tracked per condition, Mantel-Cox log-rank test: p=0.8085. (F) 150 cells tracked per condition, Mantel-Cox log-rank test: p=0.0001. (L) individual q values from all enriched terms are as follows: Reactome, from top to bottom: 0.000022, 0.000029, 0.0017.
- Figure 4: (D) n = 3 independent differentiations; values represent the mean ± standard error of the mean (SEM), Unpaired t test (two-tailed): CRYZ ***P=0.0002, HSPB1 ****P<0.0001. (G) n = 3 independent differentiations; values represent the mean ± standard error of the mean (SEM), Two-way ANOVA:

*p<0.05, **p<0.01, ***p<0.001. (H) n = 2-3 independent differentiations; values represent the mean ± standard error of the mean (SEM), Two-way ANOVA: *p<0.05. (I) 144 cells tracked per condition, Mantel-Cox log-rank test: WT vs R156X (10 μM) p<0.0001, WT vs R156X (20 μM) p<0.0001. (J) 150 cells tracked per condition, Mantel-Cox log-rank test: WT+ORF vs R156X+ORF p<0.0001, R156X+HSF1 vs R156X+ORF p<0.0001, R156X+HSF1 vs WT+HSF1 p=0.177.

## Data Availability

The mass spectrometry proteomics data have been deposited to the MassIVE repository with the identifier: (MSV000095891). Further information and requests for resources and reagents should be directed to and will be fulfilled by the Lead Contact, Evangelos Kiskinis pending scientific review and a completed material transfer agreement. Requests for these items should be submitted to.

---

**Figure S1.**
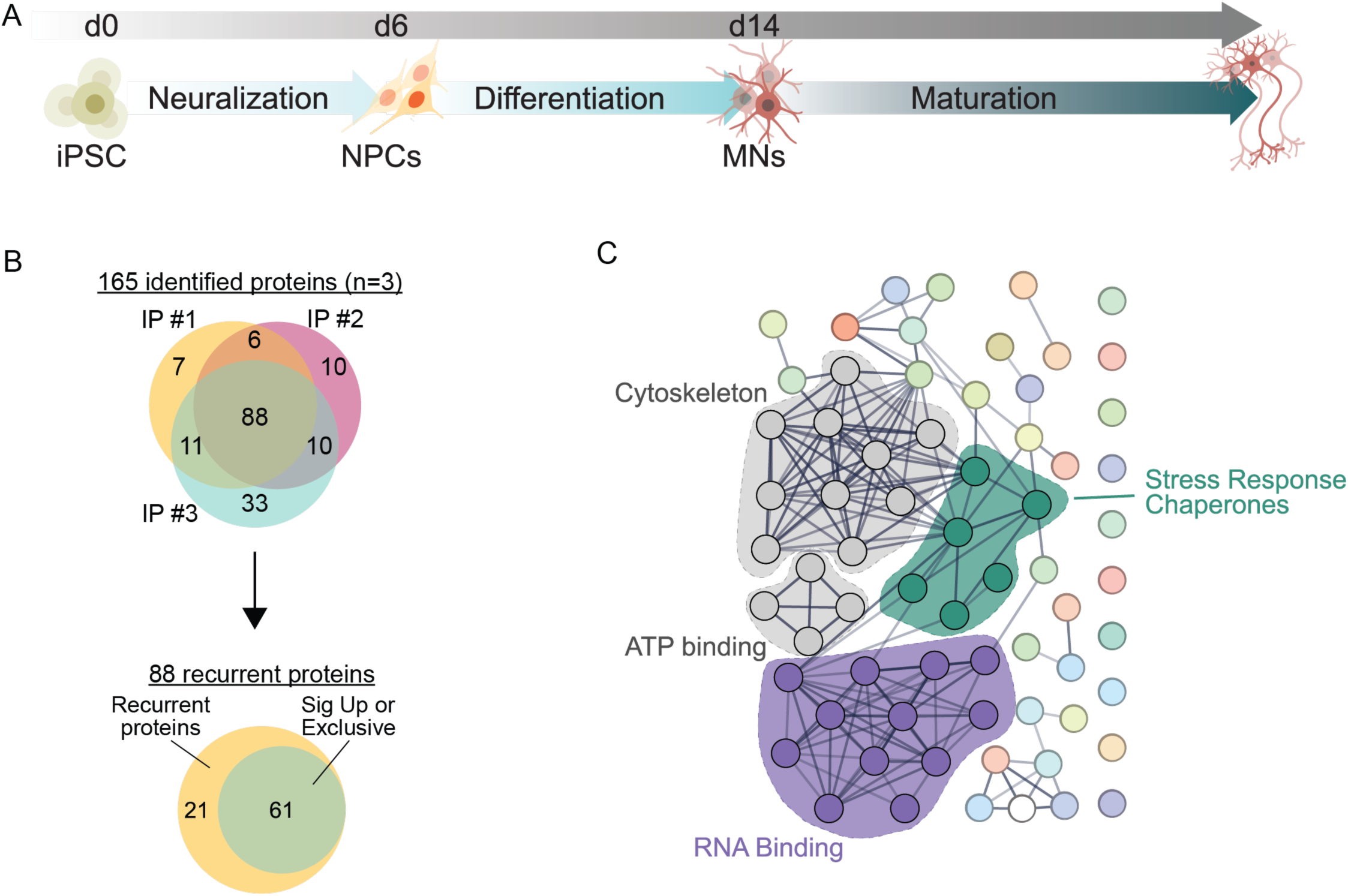
(A) **Schematic of 14-day small-molecular iPSC differentiation to lower motor neurons**. (B) Venn diagram of 88 proteins redundantly identified across n = 3 experiments. Of those 88, 61 proteins were significantly enriched in DNAJC7 IP or exclusively identified within the DNAJC7 IP. (C) STRING analysis of DNAJC7 interactome. Colors correspond to relevant identified GO enrichment pathways.

**Figure S2.**
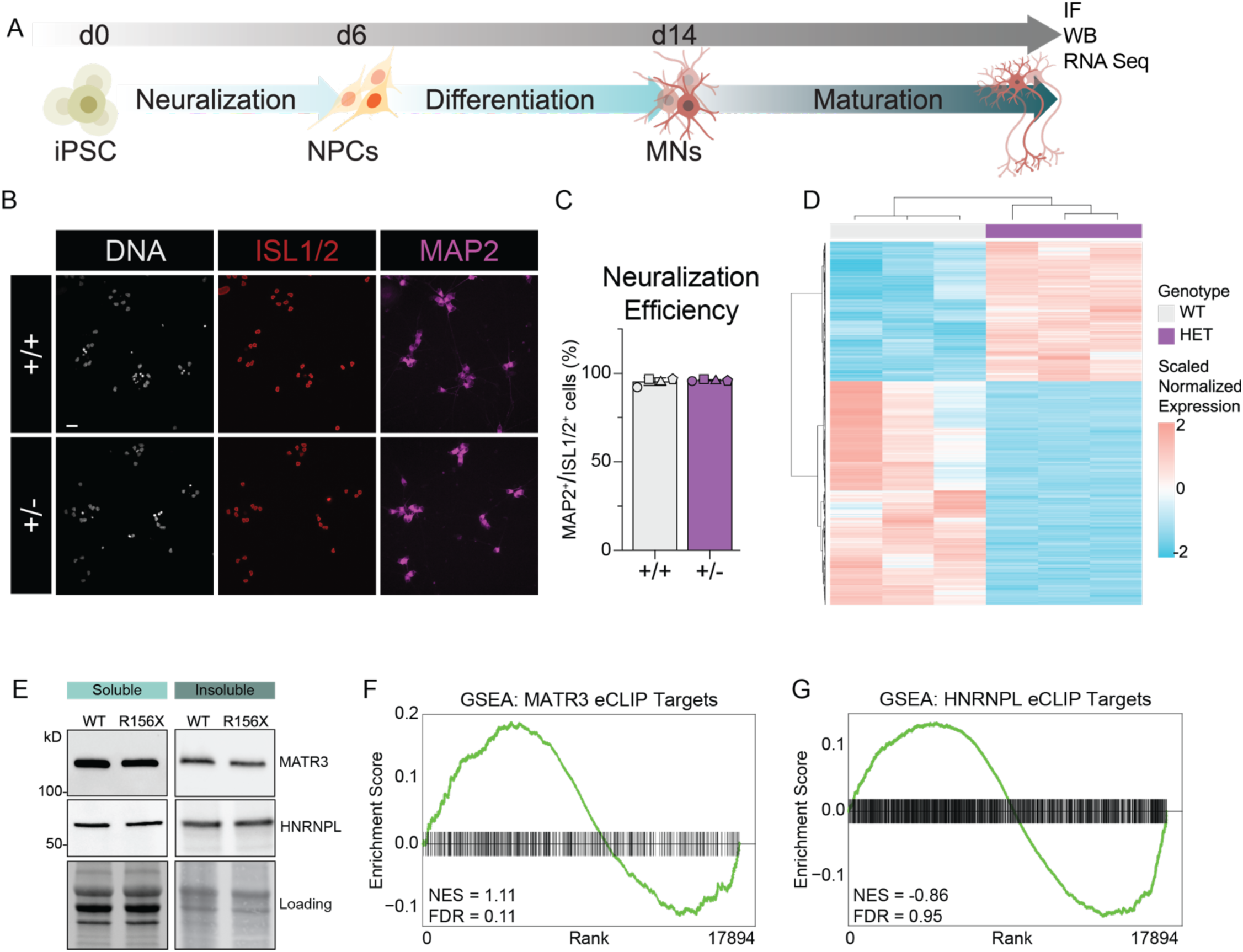
DNAJC7 haploinsufficiency disrupts HNRNPU solubility and target mRNA expression. (A) Schematic of iPSC differentiation into lower motor neurons followed by immunofluorescence (IF), WB or RNA Sequencing following maturation of 50 days in culture. (B) Confocal images of MNs derived from DNAJC7 isogenic pair immunolabeled with anti ISL1/2, anti MAP2, and Hoechst 33342. Scale bar, 25 μm. (C) Quantification of B, values represent the mean ± standard error of the mean (SEM). Experiments are represented by distinct shaped symbols. N = 4. (D) Heat map of differentially expressed genes (FDR < 0.05) in RNA Seq of R156X vs isogenic control, N = 3. (E) WB images of soluble and insoluble MATR3 and HNRNPL proteins levels from MN lysate derived from isogenic pairs. (F and G) Non-significant gene set enrichment of MATR3 and HNRNPL eCLIP targets (Kolmogorov–Smirnov test).

**Figure S3.**
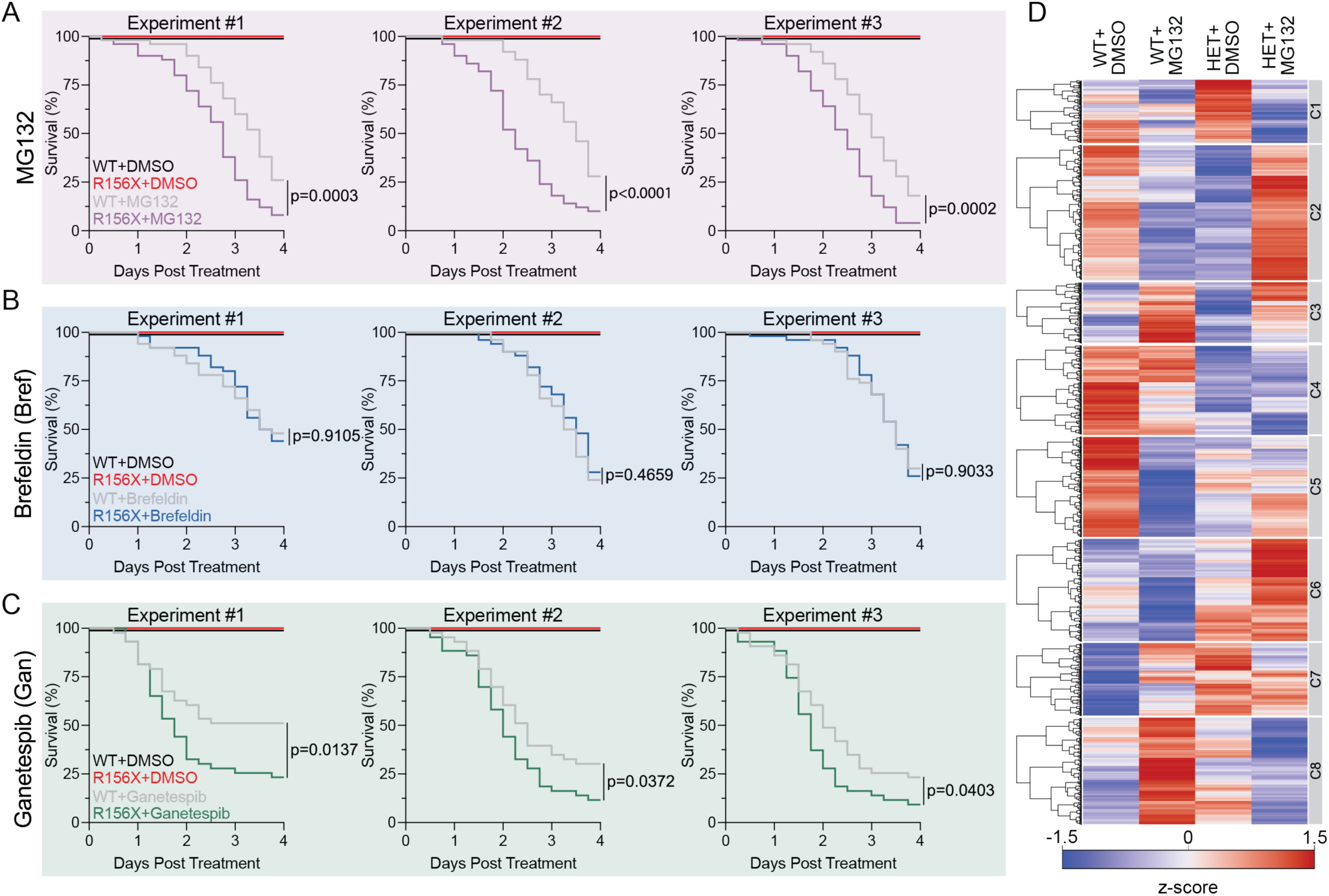
DNAJC7 haploinsufficiency sensitizes MNs to proteotoxic stress. (A) Kaplan-Meier survival curve of MNs survival following MG132 or DMSO control. Three independent differentiations separated. 50 cells per condition per experiment, Mantel-cox log-rank test: p=0.0003, p<0.0001, p=0.0002. (B) Kaplan-Meier survival curve of MNs survival following Brefeldin or DMSO control. Three independent differentiations separated. 43 cells per condition per experiment, Mantel-cox log-rank test: p=0.9105, p=0.4659, p=0.9033. (C) Kaplan-Meier survival curve of MNs survival following Ganetespib or DMSO control. Three independent differentiations separated. 50 cells per condition per experiment, Mantel-cox log-rank test: p=0.0137, p=0.0372, p=0.0403. (D) Heat map of hierarchically clustered (k-means) of average relative protein abundance for each group from TMT-MS. N = 4 independent differentiations per condition.

**Figure S4.**
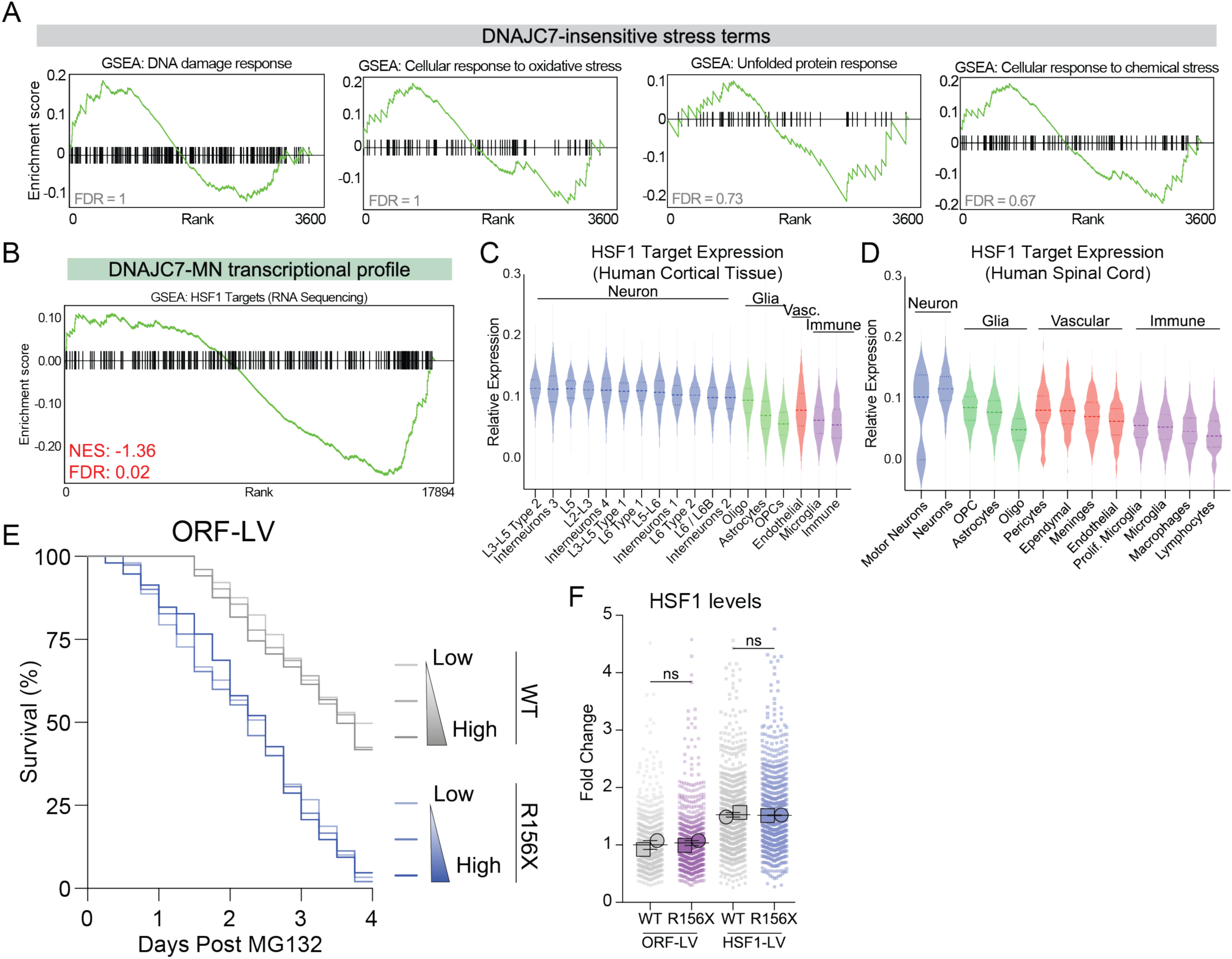
DNAJC7 regulates stress-induced activation of HSF1 and activation of HSF1 rescues the sensitivity of mutant DNAJC7-MN to stress. (A) GSEA of non-enriched “stress” related pathways in baseline proteomic dataset. DNA damage response (MSigDB#: M13636), unfolded protein response (MSigDB#: M5922), cellular response to oxidative stress (MSigDB#: M45123), and cellular response to chemical stress (MSigDB#: M29264). (B) GSEA of HSF1 transcriptional targets (MSigDB#: M19734) de-enrichment in RNA transcriptional profile of DNAJC7-MNs. Kolmogorov–Smirnov test: NES = -1.36, FDR = 0.02. (C and D) Violin plots of relative expression of HSF1 transcriptional targets (MSigDB#: M19734) as a group in human cortical (left) or spinal (right) tissue. Subtypes are categorized into Neuron, Glia, Vascular (Vasc.), or Immune cellular subgroups. (E) Kaplan-Meier survival curve of MNs survival following MG132 with ORF-LV escalating doses of ORF-LV. 36- 50 cells tracked per condition. (F) Superplots of relative expression of HSF1 following HSF1-LV or ORF-LV transduction. Experiments are represented by distinct shaped symbols, individual cell values plotted in background. N = 2, Sidak’s multiple comparisons test: WT vs R156X (ORF) p=0.9898, WT vs R156X (HSF1) p=0.9718.

**Table S1.** Results of LC-MS/MS DNAJC7 IP vs IgG IP.

| Alternate ID | Log2FC | P Value |
| --- | --- | --- |
| AASS | Inf | 0.00532813 |
| ACTC1 | 3.22706891 | 0.00185401 |
| ACTG1 | 1.88535743 | 0.00056096 |
| ACTR1A | Inf | 0.00108562 |
| ALB | -0.5986374 | 0.02876834 |
| ALK | -0.5145732 | 0.41686553 |
| ATP5F1A | Inf | 4.1619E-05 |
| ATP5F1B | Inf | 0.08508565 |
| ATP5F1C | Inf | 0.00038817 |
| ATP6V1H | Inf | 0.00749043 |
| CAND1 | Inf | 0.03994197 |
| COPA | Inf | 0.05330059 |
| CRMP1 | Inf | 0.03584222 |
| DCD | -0.6166714 | 0.34141976 |
| DCLK1 | Inf | 0.03994197 |
| DCTN2 | Inf | 0.07982533 |
| DCX | Inf | 0.00048413 |
| DNAJC7 | Inf | 0.02727733 |
| DPYSL3 | Inf | 0.10173079 |
| DSP | -0.3785116 | 0.63876821 |
| DYNC1H1 | 6.10852446 | 0.18347201 |
| EEF1A1 | 2.66675659 | 0.00420024 |
| EEF1G | Inf | 0.0013239 |
| EEF2 | Inf | 0.06384816 |
| EPPK1 | Inf | 0.00749043 |
| FASN | Inf | 0.00862393 |
| GAP43 | Inf | 0.0079662 |
| GAPDH | 0.5849625 | 0.10620776 |
| HBA1 | -0.8171359 | 0.00377202 |
| HBB | -0.5849625 | 0.070484 |
| HBG1 | -0.7484612 | 0.10639791 |
| HIST1H2BL | -1.5849625 | 0.070484 |
| HIST2H2AC | -1.0703893 | 0.10616646 |
| HNRNPC | Inf | 0.00658385 |
| HNRNPD | Inf | 0.01944177 |
| HNRNPK | Inf | 0.02696532 |
| HNRNPL | Inf | 0.00130546 |
| HNRNPU | Inf | 0.08279449 |
| HPX | -1.8073549 | 0.05831181 |
| HRNR | -0.334419 | 0.6638042 |
| HSP90AB1 | 2.12338242 | 0.00713971 |
| HSPA1A | Inf | 3.8149E-05 |
| HSPA8 | 3.66296501 | 0.0009402 |
| HUWE1 | Inf | 0.050265 |
| IGF2BP1 | 1.73696559 | 0.05723523 |
| ILF2 | Inf | 0.00749043 |
| ILF3 | Inf | 0.01007969 |
| INA | 4.7548875 | 0.00033753 |
| JUP | -0.8744691 | 0.58228482 |
| KHDRBS1 | Inf | 0.00219213 |
| KIF3A | 4.08746284 | 0.01288669 |
| KPRP | -0.6166714 | 0.58106771 |
| KRT1 | -0.0412816 | 0.88158054 |
| KRT10 | -0.180938 | 0.45181364 |
| KRT14 | -0.7399372 | 0.05336884 |
| KRT16 | -0.6764609 | 0.14050629 |
| KRT2 | -0.2642126 | 0.26694695 |
| KRT5 | -0.2715964 | 0.16378998 |
| KRT6A | -0.1554545 | 0.56959249 |
| KRT8 | Inf | 0.00053253 |
| KRT9 | -0.3520601 | 0.18475469 |
| LDHA | 0.80735492 | 0.34864114 |
| LRPPRC | Inf | 0.21999967 |
| MAP1B | Inf | 0.00388254 |
| MTHFD1 | Inf | 0.00377202 |
| NEFL | Inf | 0.0015411 |
| NEFM | Inf | 0.00121532 |
| NONO | Inf | 0.02242122 |
| PFKP | Inf | 0.11611652 |
| PKM | Inf | 0.14932117 |
| PRPH | 5.57742883 | 4.6256E-05 |
| PRSS1 | -0.3625701 | 0.72807179 |
| SFXN1 | Inf | 0.00532813 |
| SHROOM3 | -0.6100535 | 0.15183454 |
| SLC25A4 | Inf | 0.00452406 |
| SLC25A6 | Inf | 0.0009402 |
| TUBA1A | 5.07201707 | 0.00256135 |
| TUBA1B | 5.57137344 | 0.00335634 |
| TUBA4A | Inf | 0.00149455 |
| TUBB | 3.57613865 | 0.00029025 |
| TUBB2A | 3.71933992 | 0.00023504 |
| TUBB3 | 4.61667136 | 0.00057774 |
| TUBB4A | Inf | 0.00064663 |
| TUBB4B | Inf | 0.00024815 |
| TUBB6 | Inf | 0.00178996 |
| VDAC3 | Inf | 0.05723523 |
| VIM | Inf | 1.7159E-05 |
| VPS33A | Inf | 0.11340392 |

